# Existence and Localization of a Limit Cycle in a Class of Benchmark Biomolecular Oscillators

**DOI:** 10.64898/2026.04.10.717673

**Authors:** Sidhanta Mohanty, Shaunak Sen

## Abstract

Oscillatory behaviour is important in multiple biological contexts. However, the inherent nonlinearity and high dimensionality of mathematical models in biology makes proving the existence and the localization of limit cycle oscillations challenging. Here, we provided an elementary proof for the existence and a method for rigorously localizing the oscillatory solutions in a class of benchmark biomolecular oscillators. To construct the proof, we used a geometric approach based on Brouwer’s Fixed Point theorem. We constructed a toroidal-like manifold within a positively invariant set by removing the hypervolume containing the fixed point and the trajectories converging to it along its stable manifold. We showed that the vector field describing the system dynamics maps a cross section of the toroidal-like manifold onto itself. The existence of a limit cycle solution in this manifold was guaranteed by Brouwer’s Fixed Point theorem. For different sets of initial conditions in these cross-sections, we used an interval-based Reachability Analysis to localize the oscillatory behaviour that complements the Brouwer’s Fixed Point theorem approach. These results add a simple and elegant approach to demonstrating the existence of limit cycles in biomolecular systems as well as a method for rigorous localization of the region of existence.

## I. Introduction

**T**he Poincaré-Bendixson theorem provides a way to demonstrate the existence of limit cycle oscillations in two-dimensional systems [1]. The basic idea is to construct a positively invariant set without steady states so that the trajectories converge to the limit cycle. An explicit solution of the limit cycle is not needed. This theorem is not applicable in dimensions greater than two because of the possibility of more complex attractors and chaotic behaviour. Limit cycles in systems of dimension greater than two, however, are typically found in biological systems, where they underlie crucial control processes such as circadian rhythms and the cell cycle [2]. For example, the famous Goodwin oscillator has a negative feedback loop wrapping a chain of intermediate biomolecules so that the overall length of the chain is greater than or equal to three [3]–[5]. Another canonical example is a ring oscillator with an odd number (≥ 3) of genes that sequentially repress each other [6]. The experimental demonstration of the repressilator, a ring oscillator with three genes, heralded a new era for synthetic biology [7]. Therefore, methods to establish the existence and rigorous enclosure of limit cycles in higher-dimensional systems are needed.

While various approaches have been used to prove the existence of limit cycles in biological systems of dimensions greater than or equal to three, the region of existence is relatively large and conservative. Like the Poincare-Bendixson theorem, these approaches seek to establish the existence of a limit cycle without an explicit analytical solution. One approach is to analyze biological oscillators in a Boolean framework using kinetic logic models [6], [8]. A geometric approach based on Brouwer’s fixed point theorem was used to prove the existence of a limit cycle in the three-dimensional Goodwin oscillator [9], which we recently adapted for the repressilator [10]. This approach was subsequently generalized to an n-dimensional Goodwin oscillator [11]. This approach has been referred to as ‘elaborate’ [12] and ‘laborious’ [13], but provides concrete insight into the limit cycle behaviour. An n-dimensional version of the Poincare-Bendixson Theorem was developed by for monotone cyclic feedback systems [14]. The above ideas were used to address the existence and stability in ring oscillators and their variants [12], [15]. An explicit characterization of biological parameter sets for the existence of oscillations for the ring oscillators was also reported [16]–[18]. Recent work has expanded on the notion of 2-cooperativity to prove the existence of limit cycles with wide applicability in biology [19]–[21]. Typical estimates of the region of existence of the limit cycle obtained using the above approaches are conservative in nature, and rigorous tighter enclosures are desirable.

In this work, we localized the region of existence of the limit cycle in a benchmark biomolecular oscillators using the Brouwer’s Fixed Point theorem and interval-based Reachability Analysis. We provided an alternative, elementary proof for the existence of a limit cycle in cyclic gene regulatory network with odd number of genes. For the existence proof, we constructed a positively invariant set for oscillator dynamics by analyzing the direction of the vector field at the outer facets of a hypercube. We constructed a torus-like structure by removing a hypervolume associated with the unstable steady state and the trajectories approaching it along its stable manifold. We showed that a cross-section of this torus-like region maps onto itself, thus proving the existence of a limit cycle. For different sets of initial conditions on these cross-sections, we rigorously evolved the network dynamics using Interval Analysis to localize the limit cycle.

## II. Brouwer’s Fixed Point Theorem

Brouwer’s Fixed Point theorem can be used to ascertain the presence of a fixed point. The Brouwer’s Fixed Point theorem states:

### Theorem II.1

([22]). *Let S* ⊂ ℝ^*n*^ *be convex and compact, and let T*: *S* → *S be continuous. Then T has a fixed point in S*.

## III. Existence of Limit Cycle

In this section, an elementary proof for the existence of a limit cycle in *m*-gene cyclic regulatory network based on Brouwer’s Fixed Point theorem is presented.

Let *x*_1_, *x*_2_, …, *x*_*m*_ represent the proteins of the *m*-gene cyclic regulatory network. The differential equations describing the dynamics of the network are given in (1). A symmetric model with only the protein dynamics, ignoring mRNA dynamics, is chosen for simplicity.

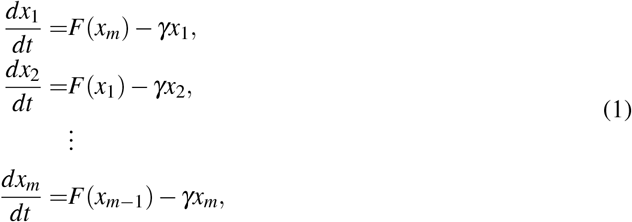

where *x_1_, x_2_*, …, *x*_*m*_ are protein concentrations. *α, γ, k*, and *n* are positive constants, 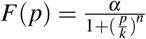 is the Hill function, *n* is the Hill coefficient, and *m* is an odd number.

The steady state of the cyclic regulatory network in (1) is unique. The uniqueness can be demonstrated graphically [2]. Denote the steady state as (*x*_10_, *x*_20_, …, *x*_*m*0_). So,

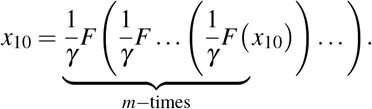

The stability of the steady state can be assessed through the eigenvalues of the Jacobian of the linearization of the system in (1) at the steady state (*x*_0_, *x*_0_, …, *x*_0_). The eigenvalues of the Jacobian are:

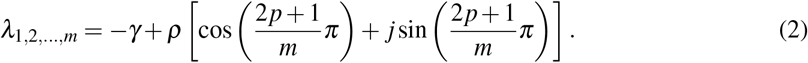

where *p* = 0, 1, …, *m* − 1 and *ρ* = −*F*^*′*^(*x*_0_). An eigenvalue is real and negative, *λ*_1_ = −(*γ* + *ρ*). The remaining eigenvalues are complex in nature. For certain values of *p*, the cosine component in the real part of the eigenvalues become negative. This implies that the corresponding eigenvalues will certainly have negative real parts. We used the instability of the steady state to prove the following sufficient condition for the existence of a limit cycle.

### Theorem III.1.

*There exists a limit cycle solution to the cyclic gene regulatory network in* (1) *if* 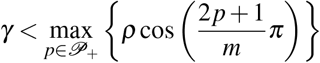, *where* 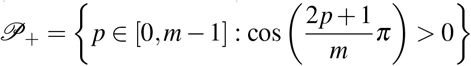.

*Proof*. For certain values of *p* in (2), the cosine component in the real part of the eigenvalues becomes negative. This implies that the corresponding eigenvalues will certainly have a negative real part. This contributes to the possible stability of the steady state. However, the instability of the steady state implies that nearby trajectories are repelled from the steady state and may converge to a closed invariant set, such as a limit cycle. To ensure that the steady state is unstable, at least one of the complex conjugate pair of eigenvalues should have a positive real part. For the complex eigenvalues to have a positive real part

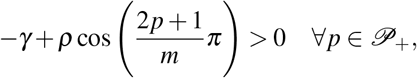

where, 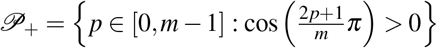. So, to ensure atleast one of the complex conjugate pair of eigenvalues have positive real part

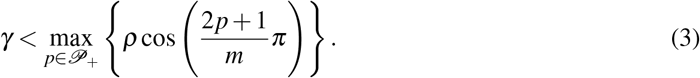

Upon establishing the instability of the steady state, a geometric approach is used to prove the existence of a limit cycle within the hypercube 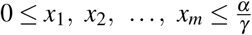.

### Construction of an Invariant Box

To demonstrate the existence of limit cycle in the *m*-gene cyclic regulatory network described by (1), we showed that the hypercube is a positively invariant set. This is done by computing the inner product between the vector field characterized by the system dynamics within the hypercube and the normal to the outer facets of the hypercube,

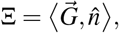

where *G* is the vector field that describes the dynamics of the *m*-gene cyclic regulatory network and 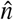 is the unit vector normal to the facet. We considered the unit normal vectors for the different facets of the hypercube directed inward. The value of Ξ determines the direction of the flow of the vector field through the facets. So,

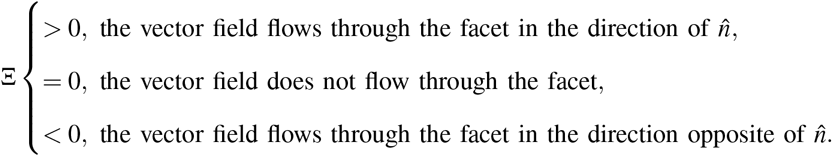

The vector field described by the system dynamics in (1) is

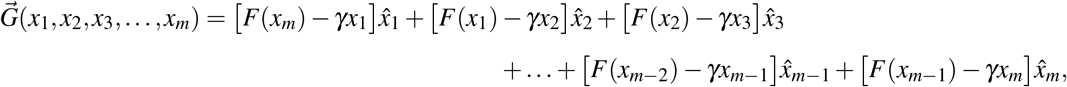

where 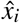 (for *i* = 1, 2, …, *m*) are the unit vectors in the direction of *x*_*i*_ axis. We calculated the inner product (Ξ) of the vector field with unit normal vector to the different facets. The normal unit vector is always considered to be directed into the hypercube. The outer facets of the hypercube are *x*_1_ = 0, 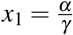, *x*_2_ = 0, 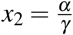, …, *x*_*m*_ = 0, and 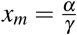. We computed the inner product between the vector field characterized by the system dynamics within the hypercube and the normal to the outer facets of the hypercube.

The inner product between the vector field characterized by the system dynamics within the hyper-cube and the normal to the outer faces of the hypercube is given in Table I. For protein concentrations between 0 and 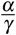, the value of Ξ is always positive. This implies that the vector field flows through the hyperplane into the hypercube. Thus, the vector field flowing through all the outer surfaces is always directed into the hypercube, making the hypercube a positively invariant box.

**Table I:**
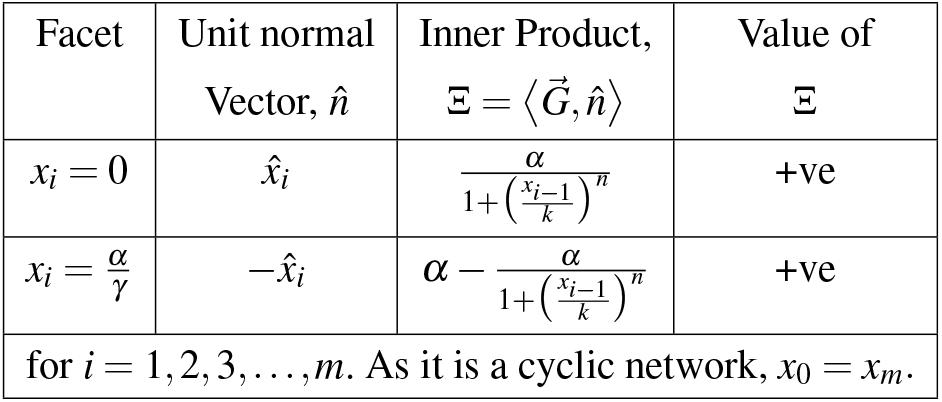
Value of the inner product between the vector field and the normal to the outer facets of the hypercube.

### Construction of an *m* - dimensional Toroidal-like Manifold

Trajectories approach the unique steady state along the path defined by the eigenvector of the negative eigenvalue. We calculated the eigenvector for the real negative eigenvalue. Let the eigenvector be 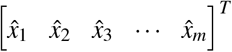. So,

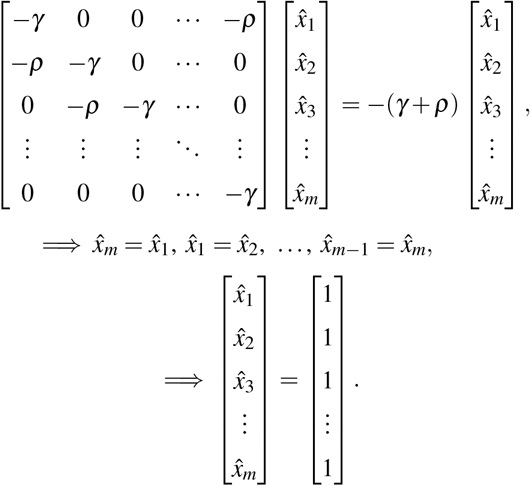

In the direction of the eigenvector 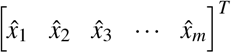, we cut out a hypervolume from the given hypercube such that the steady state and the trajectories approaching it along this singular path are eliminated. This hypervolume may be visualized as a cyclinder with a small radius *ε >* 0 along the eigenvector.

For some values of *p*, there is a possibility that some of these complex conjugate eigenvalues may have negative real part. So, we calculated the eigenvectors associated with such complex eigenvalues and removed the corresponding hypervolume along it. Let the eigenvector associated with such complex eigenvalues be 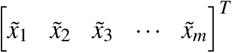. So,

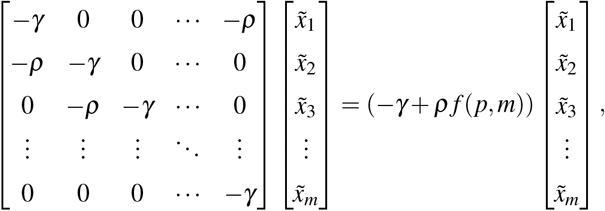

where 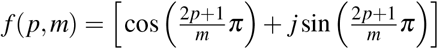 and we have

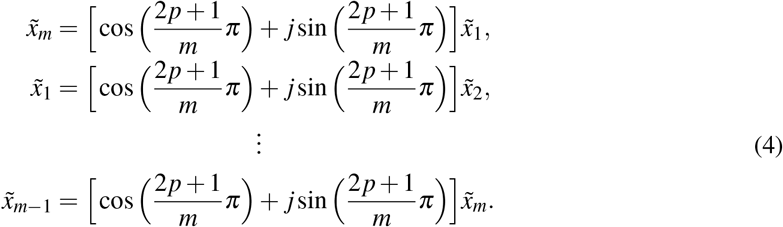

Using De Moivre’s theorem on (4), we get the eigenvector as:

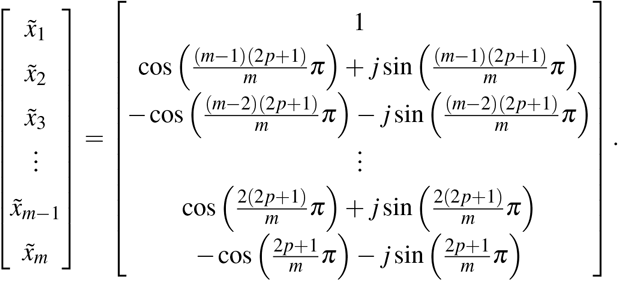

Although complex eigenvectors are not elements of the real space, their real and imaginary components reside within it. We divided the above eigenvector into its real and imaginary parts. These are the eigenvectors associated with a pair of complex conjugate eigenvalues,

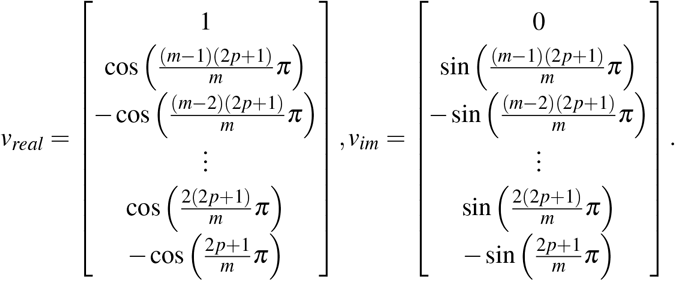

Because some of these complex conjugate eigenvalues could have a negative real part, we removed the hypervolume associated with the linear combination of *v*_*real*_ and *v*_*im*_.

So, we obtained an *m*-dimensional toroidal-like manifold without the steady state. All the trajectories that enter this manifold stay inside because the hypercube is a positively invariant set and there are not trajectories that approach the steady state.

### Continuous Map of a Cross-Section Onto Itself

We divided the hypercube into 2^*m*^ smaller hyperrectangles using the hyperplanes *x*_*i*_ = *x*_0_ (for *i* = 1, 2, 3, …, *m*). Further, we computed the flow that traverses every facet within the smaller hyperrectangles.

The flow of the vector field through the outer facets of the hypercube obtained in Table I, combined with the flow through the hyperplanes *x*_*i*_ = *x*_0_ (for *i* = 1, 2, 3, …, *m*) obtained in Table II, is used to determine the effective flow from one smaller hyperrectangle to its adjacent ones. We considered one of the smaller hyperrectangles, 0 ≤ *x*_1_ ≤ *x*_0_, 0 ≤ *x*_2_ ≤ *x*_0_, …, 0 ≤ *x*_*m*_ ≤ *x*_0_, as Box 1. From Table I, we have the flow of the vector field from the facets *x*_*i*_ = 0 (for *i* = 1, 2, 3, …, *m*) into Box 1. From Table II, we have the flow of the vector field from the hyperplanes *x*_*i*_ = *x*_0_ (for *i* = 1, 2, 3, …, *m*) out of Box 1 and into the adjacent boxes. Similarly, we calculated the effective flow of the vector field for the rest of the boxes.

**Table II:**
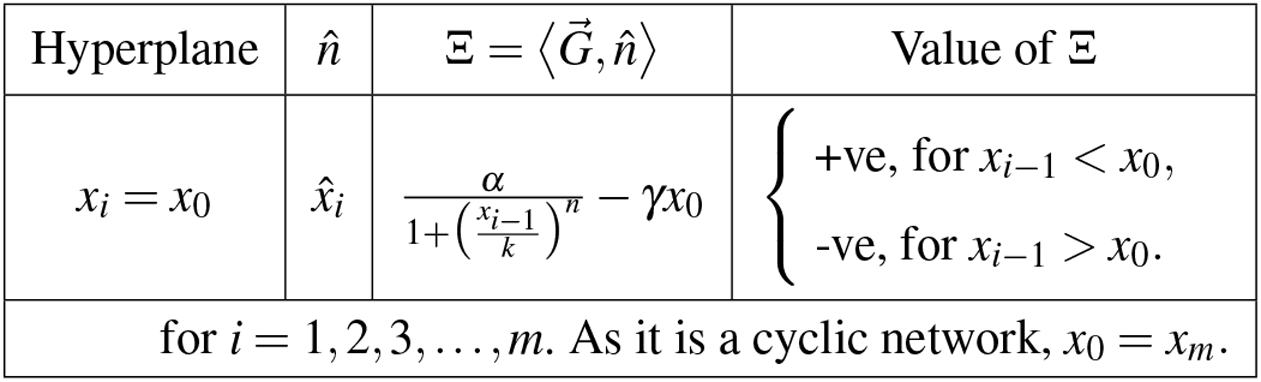
Value of the inner product between the vector field and the normal to the hyperplanes *x*_*i*_ = *x*_0_ (for *i* = 1, 2, 3, …, *m*).

Upon inspection of the effective flow from one hyperrectangle to another, we observed that the vector field forms a closed path within certain hyperrectangles. This showed that a cross section of these smaller hyperrectangles mapped onto itself. By Brouwer’s fixed point theorem, there exists a fixed point, which is a limit cycle for the system under consideration. The above construction also provides a rigorous bound on the region within the toroidal-like manifold where the limit cycle exists. □

To summarize, we obtained the region in the hypercube where the limit cycle exists. This is done by constructing a toroidal-like manifold within a positively invariant set obtained by eliminating a hypervolume containing the steady state and the trajectories approaching it. Finally, we used Brouwer’s Fixed Point theorem on a cross section of this manifold, which map onto itself, to show the existence of limit cycle.

## IV. Case Study

### A. 5-gene cyclic regulatory network

The differential equations describing the dynamics of the 5-gene cyclic regulatory network are given in (5).

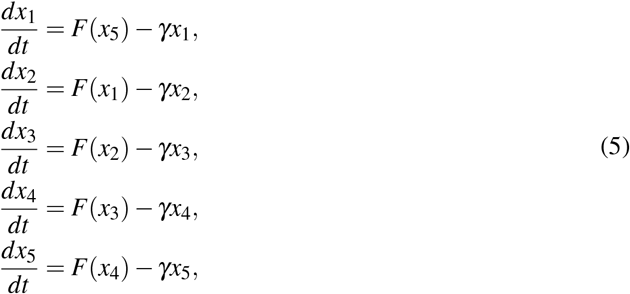

where *x*_1_, *x*_2_, *x*_3_, *x*_4_, and *x*_5_ are the protein concentrations with 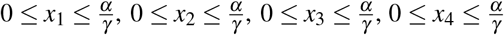, and 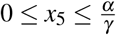. *α, γ, k*, and *n* are positive constants. 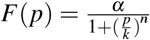 is the Hill function and *n* is the Hill coefficient. The eigenvalues of the Jacobian after the linearization of the system in (5) at the steady state (*x*_0_, *x*_0_, *x*_0_, *x*_0_, *x*_0_) is

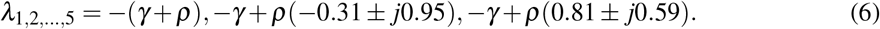

The 5-gene regulatory network has a negative real eigenvalue. Along with it, the complex conjugate eigenvalues could also have a negative real part. Guided by Theorem III.1, we illustrated the existence of the limit cycle by considering the case where one pair of the complex conjugate eigenvalues has a positive real part.

#### 1) Construction of an invariant box

We calculated the flow of the vector field through the different facets of the hypercube 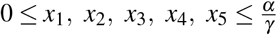. This is calculated along the unit normal vector of the various facets directed into the hypercube. Using the value of Ξ, we determined the flow of the vector field through the various facets and checked if the hypercube constituted a positively invariant set.

The vector field described by the system dynamics in (5) is

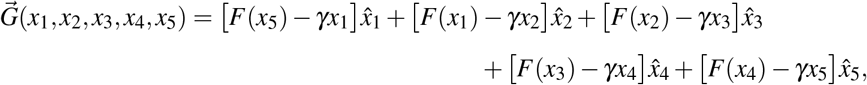

where 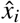 (for *i* = 1, 2, …, 5) are the unit vectors in the direction of *x*_*i*_ axis. We calculated the value of Ξ between the vector field and the unit normal vector to the different facets. The unit normal vector is always considered to be directed into the hypercube.

The value of the inner product between the vector field and the normal to the facets directed into the hypercube is given in Table III. The values of Ξ are positive for every outer facet of the hypercube. This indicated that the flow of the vector field described by the system dynamics is always directed inward and stays inside the hypercube, making it a positively invariant set.

**Table III:** Value of the inner product between the vector field and the normal to the outer facets of the hypercube.

| Facet | Unit normal<br>Vector, $\hat{n}$ | Inner Product,<br>$\Xi = \langle \vec{G}, \hat{n} \rangle$ | Value of<br>$\Xi$ |
| --- | --- | --- | --- |
| $x_1 = 0$ | $\hat{x}_1$ | $\frac{\alpha}{1 + \left(\frac{x_5}{k}\right)^n}$ | +ve |
| $x_1 = \frac{\alpha}{\gamma}$ | $-\hat{x}_1$ | $\alpha - \frac{\alpha}{1 + \left(\frac{x_5}{k}\right)^n}$ | +ve |
| $x_2 = 0$ | $\hat{x}_2$ | $\frac{\alpha}{1 + \left(\frac{x_1}{k}\right)^n}$ | +ve |
| $x_2 = \frac{\alpha}{\gamma}$ | $-\hat{x}_2$ | $\alpha - \frac{\alpha}{1 + \left(\frac{x_1}{k}\right)^n}$ | +ve |
| $x_3 = 0$ | $\hat{x}_3$ | $\frac{\alpha}{1 + \left(\frac{x_2}{k}\right)^n}$ | +ve |
| $x_3 = \frac{\alpha}{\gamma}$ | $-\hat{x}_3$ | $\alpha - \frac{\alpha}{1 + \left(\frac{x_2}{k}\right)^n}$ | +ve |
| $x_4 = 0$ | $\hat{x}_4$ | $\frac{\alpha}{1 + \left(\frac{x_3}{k}\right)^n}$ | +ve |
| $x_4 = \frac{\alpha}{\gamma}$ | $-\hat{x}_4$ | $\alpha - \frac{\alpha}{1 + \left(\frac{x_3}{k}\right)^n}$ | +ve |
| $x_5 = 0$ | $\hat{x}_5$ | $\frac{\alpha}{1 + \left(\frac{x_4}{k}\right)^n}$ | +ve |
| $x_5 = \frac{\alpha}{\gamma}$ | $-\hat{x}_5$ | $\alpha - \frac{\alpha}{1 + \left(\frac{x_4}{k}\right)^n}$ | +ve |

#### 2) Construction of a 5− dimensional Toroidal-like Manifold

The existence of a unique steady state in the given hypercube indicated that certain trajectories approach the steady state along the singular path defined by the eigenvector of the negative eigenvalue. So, we calculated the eigenvector for the real negative eigenvalue. Let the eigenvector be 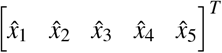. So,

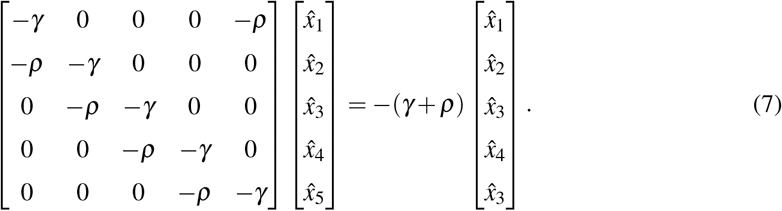

From (7), the eigenvector is found to be

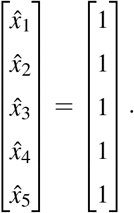

In the direction of the eigenvector 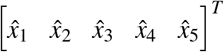, we cut out a hypervolume from the given hypercube, such that the steady state and the trajectories approaching it along the singular path are eliminated.

The regulatory network also has a pair of complex conjugate eigenvalues with negative real part. So, we calculated the eigenvectors associated with these complex eigenvalues and remove a hypervolume along it. Let the eigenvector associated with one of these complex eigenvalues be 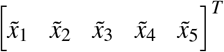. So,

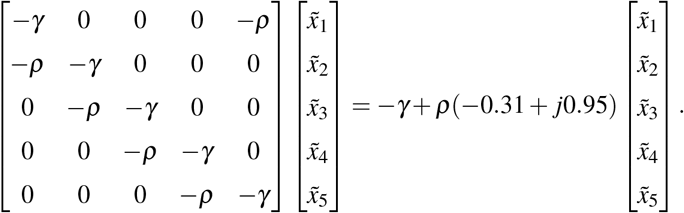

Using the relation in (4) and subsequently using the De Moivre’s theorem, we get the eigenvector as:

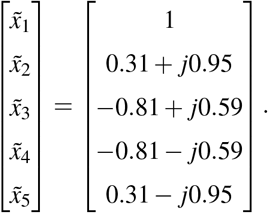

We divide the above eigenvector into its real and imaginary parts. These are the eigenvectors associated complex eigenvalue with negative real part.

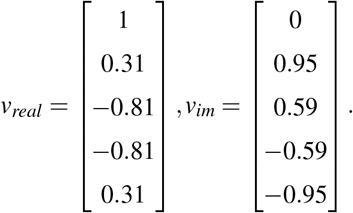

The complex conjugate eigenvalues produce conjugate eigenvectors. So, the *v*_*real*_ remains same for the complex conjugate pair. *v*_*im*_ for the pair of eigenvalues is a conjugate pair. However, the conjugate gives the same eigenvector but in the opposite direction. In the sense of removing a hypervolume, the *v*_*im*_ can be considered to be same. So, we also removed the hypervolume along the eigenvectors *v*_*real*_ and *v*_*im*_.

So, we obtained a toroidal-like manifold with no steady state. All the trajectories are inside the torus-like structure (because the hypercube is a positively invariant set).

#### 3) Continuous Map of a Cross-Section Onto Itself

The hypercube is divided into 2^5^ smaller hyperrectangles by using the hyperplanes *x*_1_ = *x*_0_, *x*_2_ = *x*_0_, *x*_3_ = *x*_0_, *x*_4_ = *x*_0_, and *x*_5_ = *x*_0_. We calculated the value of Ξ between the vector field describing the system dynamics and the unit normal vector through these hyperplanes.

The value of the inner product between the vector field and the normal to the hyperplanes *x*_1_ = *x*_0_, *x*_2_ = *x*_0_, *x*_3_ = *x*_0_, *x*_4_ = *x*_0_, and *x*_5_ = *x*_0_ is given in Table IV. Using the values of Ξ in Table III and Table IV, we determined the direction of flow of the vector field within the smaller boxes. This is summarized in Table V.

**Table IV:** Value of the inner product between the vector field and the normal to the hyperplanes *x*_*i*_ = *x*_0_ (for *i* = 1, 2, …, 5).

| Hyperplane | Unit normal Vector, $\hat{n}$ | Inner Product, $\Xi = \langle \vec{G}, \hat{n} \rangle$ | Value of $\Xi$ |
| --- | --- | --- | --- |
| $x_1 = x_0$ | $\hat{x}_1$ | $\frac{\alpha}{1 + \left(\frac{x_5}{k}\right)^n} - \gamma x_0$ | $\begin{cases} +\text{ve, for } x_5 < x_0, \\ -\text{ve, for } x_5 > x_0. \end{cases}$ |
| $x_2 = x_0$ | $\hat{x}_2$ | $\frac{\alpha}{1 + \left(\frac{x_1}{k}\right)^n} - \gamma x_0$ | $\begin{cases} +\text{ve, for } x_1 < x_0, \\ -\text{ve, for } x_1 > x_0. \end{cases}$ |
| $x_3 = x_0$ | $\hat{x}_3$ | $\frac{\alpha}{1 + \left(\frac{x_2}{k}\right)^n} - \gamma x_0$ | $\begin{cases} +\text{ve, for } x_2 < x_0, \\ -\text{ve, for } x_2 > x_0. \end{cases}$ |
| $x_4 = x_0$ | $\hat{x}_4$ | $\frac{\alpha}{1 + \left(\frac{x_3}{k}\right)^n} - \gamma x_0$ | $\begin{cases} +\text{ve, for } x_3 < x_0, \\ -\text{ve, for } x_3 > x_0. \end{cases}$ |
| $x_5 = x_0$ | $\hat{x}_5$ | $\frac{\alpha}{1 + \left(\frac{x_4}{k}\right)^n} - \gamma x_0$ | $\begin{cases} +\text{ve, for } x_4 < x_0, \\ -\text{ve, for } x_4 > x_0. \end{cases}$ |

**Table V:** Direction of flow of vector field within the smaller boxes.

| Box | Coordinates of the hyperrectangle | Direction of the flow of vector field |
| --- | --- | --- |
| 1 | $0 \leq x_1 \leq x_0$<br>$0 \leq x_2 \leq x_0$<br>$0 \leq x_3 \leq x_0$<br>$0 \leq x_4 \leq x_0$<br>$0 \leq x_5 \leq x_0$ | Box 2<br>↑<br>Box 32 ← Box 1 → Box 29<br>$\swarrow \quad \searrow$<br>Box 31 Box 32 |
| 2 | $x_0 \leq x_1 \leq \frac{\alpha}{\gamma}$<br>$0 \leq x_2 \leq x_0$<br>$0 \leq x_3 \leq x_0$<br>$0 \leq x_4 \leq x_0$<br>$0 \leq x_5 \leq x_0$ | Box 25<br>↗<br>Box 2 → Box 26<br>↘<br>Box 28 |
| 3 | $x_0 \leq x_1 \leq \frac{\alpha}{\gamma}$<br>$x_0 \leq x_2 \leq \frac{\alpha}{\gamma}$<br>$0 \leq x_3 \leq x_0$<br>$0 \leq x_4 \leq x_0$<br>$0 \leq x_5 \leq x_0$ | Box 2<br>↗<br>Box 3 → Box 18<br>↘<br>Box 27 |
| 4 | $x_0 \leq x_1 \leq \frac{\alpha}{\gamma}$<br>$x_0 \leq x_2 \leq \frac{\alpha}{\gamma}$<br>$x_0 \leq x_3 \leq \frac{\alpha}{\gamma}$<br>$0 \leq x_4 \leq x_0$<br>$0 \leq x_5 \leq x_0$ | Box 10<br>↗<br>Box 4 → Box 3<br>↘<br>Box 28 |
| 5 | $x_0 \leq x_1 \leq \frac{\alpha}{\gamma}$<br>$x_0 \leq x_2 \leq \frac{\alpha}{\gamma}$<br>$x_0 \leq x_3 \leq \frac{\alpha}{\gamma}$<br>$x_0 \leq x_4 \leq \frac{\alpha}{\gamma}$<br>$0 \leq x_5 \leq x_0$ | Box 4<br>↗<br>Box 5 → Box 17<br>↘<br>Box 19 |
| 6 | $x_0 \leq x_1 \leq \frac{\alpha}{\gamma}$<br>$x_0 \leq x_2 \leq \frac{\alpha}{\gamma}$<br>$x_0 \leq x_3 \leq \frac{\alpha}{\gamma}$<br>$x_0 \leq x_4 \leq \frac{\alpha}{\gamma}$<br>$x_0 \leq x_5 \leq \frac{\alpha}{\gamma}$ | Box 5<br>↑<br>Box 10 ← Box 6 → Box 7<br>$\swarrow \quad \searrow$<br>Box 9 Box 8 |
| 7 | $0 \leq x_1 \leq x_0$<br>$x_0 \leq x_2 \leq \frac{\alpha}{\gamma}$<br>$x_0 \leq x_3 \leq \frac{\alpha}{\gamma}$<br>$x_0 \leq x_4 \leq \frac{\alpha}{\gamma}$<br>$x_0 \leq x_5 \leq \frac{\alpha}{\gamma}$ | Box 12<br>↗<br>Box 7 → Box 13<br>↘<br>Box 14 |
| Continued on the next page. |  |  |

| Continuation of Table V ... |  |  |
| --- | --- | --- |
| Box | Coordinates of the hyperrectangle | Direction of the flow of vector field |
| 8 | $x_0 \leq x_1 \leq \frac{\alpha}{\gamma}$<br>$0 \leq x_2 \leq x_0$<br>$x_0 \leq x_3 \leq \frac{\alpha}{\gamma}$<br>$x_0 \leq x_4 \leq \frac{\alpha}{\gamma}$<br>$x_0 \leq x_5 \leq \frac{\alpha}{\gamma}$ | Box 11<br>↗<br>Box 8 → Box 16<br>↘<br>Box 17 |
| 9 | $x_0 \leq x_1 \leq \frac{\alpha}{\gamma}$<br>$x_0 \leq x_2 \leq \frac{\alpha}{\gamma}$<br>$0 \leq x_3 \leq x_0$<br>$x_0 \leq x_4 \leq \frac{\alpha}{\gamma}$<br>$x_0 \leq x_5 \leq \frac{\alpha}{\gamma}$ | Box 12<br>↗<br>Box 9 → Box 15<br>↘<br>Box 19 |
| 10 | $x_0 \leq x_1 \leq \frac{\alpha}{\gamma}$<br>$x_0 \leq x_2 \leq \frac{\alpha}{\gamma}$<br>$x_0 \leq x_3 \leq \frac{\alpha}{\gamma}$<br>$0 \leq x_4 \leq x_0$<br>$x_0 \leq x_5 \leq \frac{\alpha}{\gamma}$ | Box 13<br>↗<br>Box 10 → Box 16<br>↘<br>Box 18 |
| 11 | $0 \leq x_1 \leq x_0$<br>$0 \leq x_2 \leq x_0$<br>$x_0 \leq x_3 \leq \frac{\alpha}{\gamma}$<br>$x_0 \leq x_4 \leq \frac{\alpha}{\gamma}$<br>$x_0 \leq x_5 \leq \frac{\alpha}{\gamma}$ | Box 7<br>↗<br>Box 11 → Box 21<br>↘<br>Box 22 |
| 12 | $0 \leq x_1 \leq x_0$<br>$x_0 \leq x_2 \leq \frac{\alpha}{\gamma}$<br>$0 \leq x_3 \leq x_0$<br>$x_0 \leq x_4 \leq \frac{\alpha}{\gamma}$<br>$x_0 \leq x_5 \leq \frac{\alpha}{\gamma}$ | Box 12 → Box 24 |
| 13 | $0 \leq x_1 \leq x_0$<br>$x_0 \leq x_2 \leq \frac{\alpha}{\gamma}$<br>$x_0 \leq x_3 \leq \frac{\alpha}{\gamma}$<br>$0 \leq x_4 \leq x_0$<br>$x_0 \leq x_5 \leq \frac{\alpha}{\gamma}$ | Box 13 → Box 23 |
| 14 | $0 \leq x_1 \leq x_0$<br>$x_0 \leq x_2 \leq \frac{\alpha}{\gamma}$<br>$x_0 \leq x_3 \leq \frac{\alpha}{\gamma}$<br>$x_0 \leq x_4 \leq \frac{\alpha}{\gamma}$<br>$0 \leq x_5 \leq x_0$ | Box 5<br>↗<br>Box 14 → Box 24<br>↘<br>Box 27 |
| Continued on the next page. |  |  |
| Box | Coordinates of the hyperrectangle | Direction of the flow of vector field |
| 15 | $x_0 \leq x_1 \leq \frac{\alpha}{\gamma}$<br>$0 \leq x_2 \leq x_0$<br>$0 \leq x_3 \leq x_0$<br>$x_0 \leq x_4 \leq \frac{\alpha}{\gamma}$<br>$x_0 \leq x_5 \leq \frac{\alpha}{\gamma}$ | Box 8<br>↗<br>Box 15 → Box 20<br>↘<br>Box 26 |
| 16 | $x_0 \leq x_1 \leq \frac{\alpha}{\gamma}$<br>$0 \leq x_2 \leq x_0$<br>$x_0 \leq x_3 \leq \frac{\alpha}{\gamma}$<br>$0 \leq x_4 \leq x_0$<br>$x_0 \leq x_5 \leq \frac{\alpha}{\gamma}$ | Box 16 → Box 21 |
| 17 | $x_0 \leq x_1 \leq \frac{\alpha}{\gamma}$<br>$0 \leq x_2 \leq x_0$<br>$x_0 \leq x_3 \leq \frac{\alpha}{\gamma}$<br>$x_0 \leq x_4 \leq \frac{\alpha}{\gamma}$<br>$0 \leq x_5 \leq x_0$ | Box 17 → Box 28 |
| 18 | $x_0 \leq x_1 \leq \frac{\alpha}{\gamma}$<br>$x_0 \leq x_2 \leq \frac{\alpha}{\gamma}$<br>$0 \leq x_3 \leq x_0$<br>$0 \leq x_4 \leq x_0$<br>$x_0 \leq x_5 \leq \frac{\alpha}{\gamma}$ | Box 9<br>↗<br>Box 15 → Box 23<br>↘<br>Box 25 |
| 19 | $x_0 \leq x_1 \leq \frac{\alpha}{\gamma}$<br>$x_0 \leq x_2 \leq \frac{\alpha}{\gamma}$<br>$0 \leq x_3 \leq x_0$<br>$x_0 \leq x_4 \leq \frac{\alpha}{\gamma}$<br>$0 \leq x_5 \leq x_0$ | Box 19 → Box 26 |
| 20 | $0 \leq x_1 \leq x_0$<br>$0 \leq x_2 \leq x_0$<br>$0 \leq x_3 \leq x_0$<br>$x_0 \leq x_4 \leq \frac{\alpha}{\gamma}$<br>$x_0 \leq x_5 \leq \frac{\alpha}{\gamma}$ | Box 11<br>↗<br>Box 20 → Box 12<br>↘<br>Box 30 |
| 21 | $0 \leq x_1 \leq x_0$<br>$0 \leq x_2 \leq x_0$<br>$x_0 \leq x_3 \leq \frac{\alpha}{\gamma}$<br>$0 \leq x_4 \leq x_0$<br>$x_0 \leq x_5 \leq \frac{\alpha}{\gamma}$ | Box 21 → Box 13 |
| Continued on the next page. |  |  |
| Box | Coordinates of the hyperrectangle | Direction of the flow of vector field |
| 22 | $0 \leq x_1 \leq x_0$<br>$0 \leq x_2 \leq x_0$<br>$x_0 \leq x_3 \leq \frac{\alpha}{\gamma}$<br>$x_0 \leq x_4 \leq \frac{\alpha}{\gamma}$<br>$0 \leq x_5 \leq x_0$ | Box 14<br>↗<br>Box 22 → Box 17<br>↘<br>Box 31 |
| 23 | $0 \leq x_1 \leq x_0$<br>$x_0 \leq x_2 \leq \frac{\alpha}{\gamma}$<br>$0 \leq x_3 \leq x_0$<br>$0 \leq x_4 \leq x_0$<br>$x_0 \leq x_5 \leq \frac{\alpha}{\gamma}$ | Box 23 → Box 12 |
| 24 | $0 \leq x_1 \leq x_0$<br>$x_0 \leq x_2 \leq \frac{\alpha}{\gamma}$<br>$0 \leq x_3 \leq x_0$<br>$x_0 \leq x_4 \leq \frac{\alpha}{\gamma}$<br>$0 \leq x_5 \leq x_0$ | Box 24 → Box 19 |
| 25 | $x_0 \leq x_1 \leq \frac{\alpha}{\gamma}$<br>$0 \leq x_2 \leq x_0$<br>$0 \leq x_3 \leq x_0$<br>$0 \leq x_4 \leq x_0$<br>$x_0 \leq x_5 \leq \frac{\alpha}{\gamma}$ | Box 15<br>↗<br>Box 25 → Box 16<br>↘<br>Box 29 |
| 26 | $x_0 \leq x_1 \leq \frac{\alpha}{\gamma}$<br>$0 \leq x_2 \leq x_0$<br>$0 \leq x_3 \leq x_0$<br>$x_0 \leq x_4 \leq \frac{\alpha}{\gamma}$<br>$0 \leq x_5 \leq x_0$ | Box 26 → Box 17 |
| 27 | $0 \leq x_1 \leq x_0$<br>$x_0 \leq x_2 \leq \frac{\alpha}{\gamma}$<br>$x_0 \leq x_3 \leq \frac{\alpha}{\gamma}$<br>$0 \leq x_4 \leq x_0$<br>$0 \leq x_5 \leq x_0$ | Box 4<br>↗<br>Box 27 → Box 13<br>↘<br>Box 32 |
| 28 | $x_0 \leq x_1 \leq \frac{\alpha}{\gamma}$<br>$0 \leq x_2 \leq x_0$<br>$x_0 \leq x_3 \leq \frac{\alpha}{\gamma}$<br>$0 \leq x_4 \leq x_0$<br>$0 \leq x_5 \leq x_0$ | Box 28 → Box 16 |
| Continued on the next page. |  |  |
| Box | Coordinates of the hyperrectangle | Direction of the flow of vector field |
| 29 | $0 \leq x_1 \leq x_0$ | Box 20 |
| | $0 \leq x_2 \leq x_0$ | ↗ |
| | $0 \leq x_3 \leq x_0$ | Box 29 → Box 21 |
| | $0 \leq x_4 \leq x_0$ | ↘ |
| | $x_0 \leq x_5 \leq \frac{\alpha}{\gamma}$ | Box 23 |
| 30 | $0 \leq x_1 \leq x_0$ | Box 22 |
| | $0 \leq x_2 \leq x_0$ | ↗ |
| | $0 \leq x_3 \leq x_0$ | Box 30 → Box 24 |
| | $x_0 \leq x_4 \leq \frac{\alpha}{\gamma}$ | ↘ |
| | $0 \leq x_5 \leq x_0$ | Box 26 |
| 31 | $0 \leq x_1 \leq x_0$ | Box 21 |
| | $0 \leq x_2 \leq x_0$ | ↗ |
| | $x_0 \leq x_3 \leq \frac{\alpha}{\gamma}$ | Box 31 → Box 27 |
| | $0 \leq x_4 \leq x_0$ | ↘ |
| | $0 \leq x_5 \leq x_0$ | Box 28 |
| 32 | $0 \leq x_1 \leq x_0$ | Box 3 |
| | $x_0 \leq x_2 \leq \frac{\alpha}{\gamma}$ | ↗ |
| | $0 \leq x_3 \leq x_0$ | Box 32 → Box 23 |
| | $0 \leq x_4 \leq x_0$ | ↘ |
| | $0 \leq x_5 \leq x_0$ | Box 24 |

Upon inspection of Table V, we find that the vector field formed a continuous map of a cross section onto itself. The continuous map is as follows:

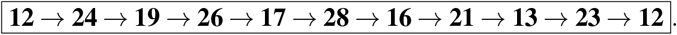

We obtained a toroidal-like manifold (which is a positively invariant set) and a cross section of this structure formed a continuous map onto itself. So, by Brouwer’s Fixed Point theorem, there exists a limit cycle.

## V. Localization of Limit Cycle

The Brouwer’s Fixed Point theorem guarantees the presence of a fixed point and thus a periodic orbit through a cross-section that maps onto itself. However, where the limit cycle trajectory lies is still undetermined. Thus localization is the natural next step in rigorously refining the region of existence of the limit cycle. We now use an interval-based Reachability Analysis to localize the region of existence. This method is based on rigorously solving the network dynamics using Interval Analysis. We consider a set of initial conditions within the region of existence that we want to localize. By the definition of limit cycle, the sets obtained using Reachability Analysis that do not map onto themselves, under the flow of the system dynamics, does not contain a limit cycle. Therefore the limit cycle can be localized by combining the existence result from the Brouwer’s Fixed Point theorem with the non-existence criterion implemented by the set-based Reachability Analysis.

This interval-based Reachability Analysis approach thus serves as a complement of the Brouwer’s Fixed Point theorem’s topological arguments using validated numerical techniques. In this approach, the geometric confinement test extends the theoretical proof by connecting the existence result to a rigorous characterization of the system’s oscillatory dynamics.

### A. Methodology

The methodology involves dividing the search space into smaller hyperrectangles and evolving the system dynamics from each region using interval-based reachability analysis to generate a flowpipe representation. For every initial box *X*_0_, the computed flowpipe is examined to detect whether the reachable sets return back to (or intersect with) the initial region. Based on this return behavior, each region is classified as: (i) Does not contain the limit cycle, (ii) May contain the limit cycle, (iii) Contains the limit cycle, or (iv) Failed boxes.

This classification is performed using the return-index detection routine `intersection`, which identifies time indices where the reachable set overlaps with the initial box *X*_0_. If no such return indices exist, the region is classified as “Does not contain the limit cycle”. To evaluate whether the return behavior is consistent with a limit cycle being contained inside the region, a geometric containment check is applied on the reachable sets. Specifically, the initial reachable set is defined as

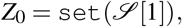

and it is compared against all return candidates

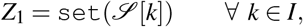

where *I* is the return index set produced by the `intersection` function. A subset test `issubset`(*Z*_1_, *Z*_0_) is then used to determine whether the return set is enclosed within the initial reachable set approximation.

The final classification is performed as follows:

- **Does not contain the limit cycle** is assigned when no return indices are detected, meaning the flowpipe never intersects the initial region.
- **May contain the limit cycle** is assigned when return indices are detected, but none of the corresponding reachable sets satisfy the subset condition with respect to *Z*_0_, indicating that the trajectory returns to the initial region but the enclosure property fails.
- **Contains the limit cycle** is assigned when all return indices detected fulfill the subset condition `issubset`(*Z*_1_, *Z*_0_).
- **Failed boxes** are recorded when reachability computation fails due to solver breakdown or numerical over approximation, in which case no reliable classification can be made.

This approach avoids expensive direct geometric comparisons over the entire flowpipe and instead relies on return detection combined with subset verification, enabling limit cycle localization over a search space.

### B. Algorithms^†^

An external loop initially calls the function `divide` to partition the entire search space into smaller hyperrectangular subsets. Each subset is then analyzed sequentially, enabling classification of the phase space and localization of regions that may contain the limit cycle of the repressilator model. For each partition *X*_0_, the reachability flowpipe is computed using the subroutine `solve_ivp`, which propagates the interval initial condition set forward in time and generates an overapproximated reachable set sequence 𝒮. If the solver fails or produces invalid bounds, the corresponding region is marked as a failed box and excluded from further analysis. If the flowpipe is successfully generated, the function `intersection` is used to identify thee indices at which the reachable set intersects the original subset *X*_0_. These indices represent the return candidates. If no such return indices are obtained, the subset is classified as “Does not contain the limit cycle”.

Subsequently, a geometric containment verification is performed using a zonotope subset test. The initial reachable set *Z*_0_ = `set`(𝒮[1]) is compared against each return candidate *Z*_1_ = `set`(𝒮[*k*]), where *k* belongs to the intersection index set. The classification decision is made based on the subset condition `issubset`(*Z*_1_, *Z*_0_), which determines whether the return sets are enclosed within the initial approximation. This hierarchical structure allows categorization of the search space into regions (i) Does not contain the limit cycle, (ii) May contain the limit cycle, or (iii) Contains the limit cycle. The box containing the steady state is removed from the reachability analysis.

The algorithm outlined below defines the process for identifying and categorizing and thereby localizing limit cycle trajectory within the phase space of the repressilator.

#### Algorithm 1: Reachability Analysis for Localization of Limit Cycle

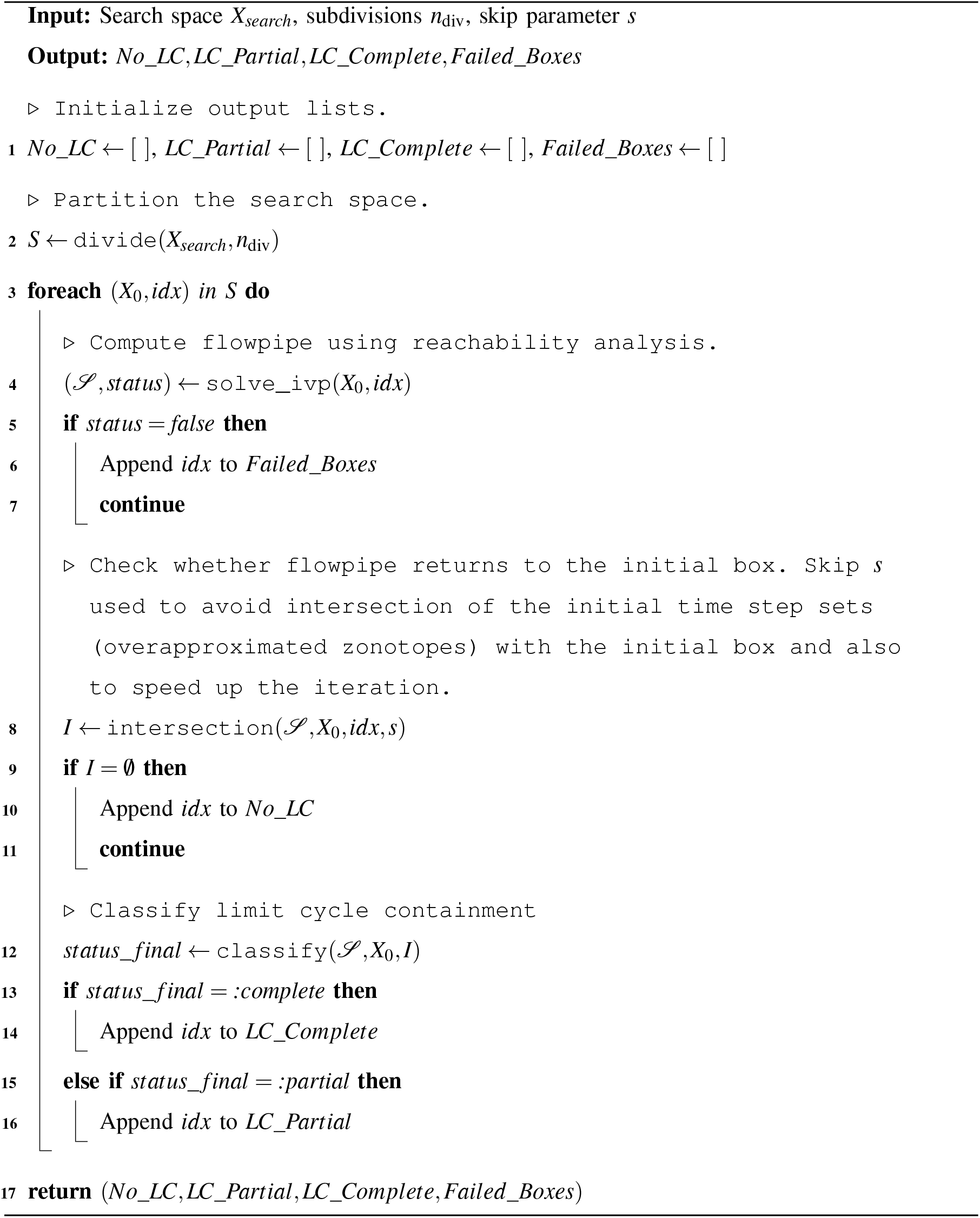

#### Algorithm 2: Subroutines for Reachability Analysis Based Localization of Limit Cycle

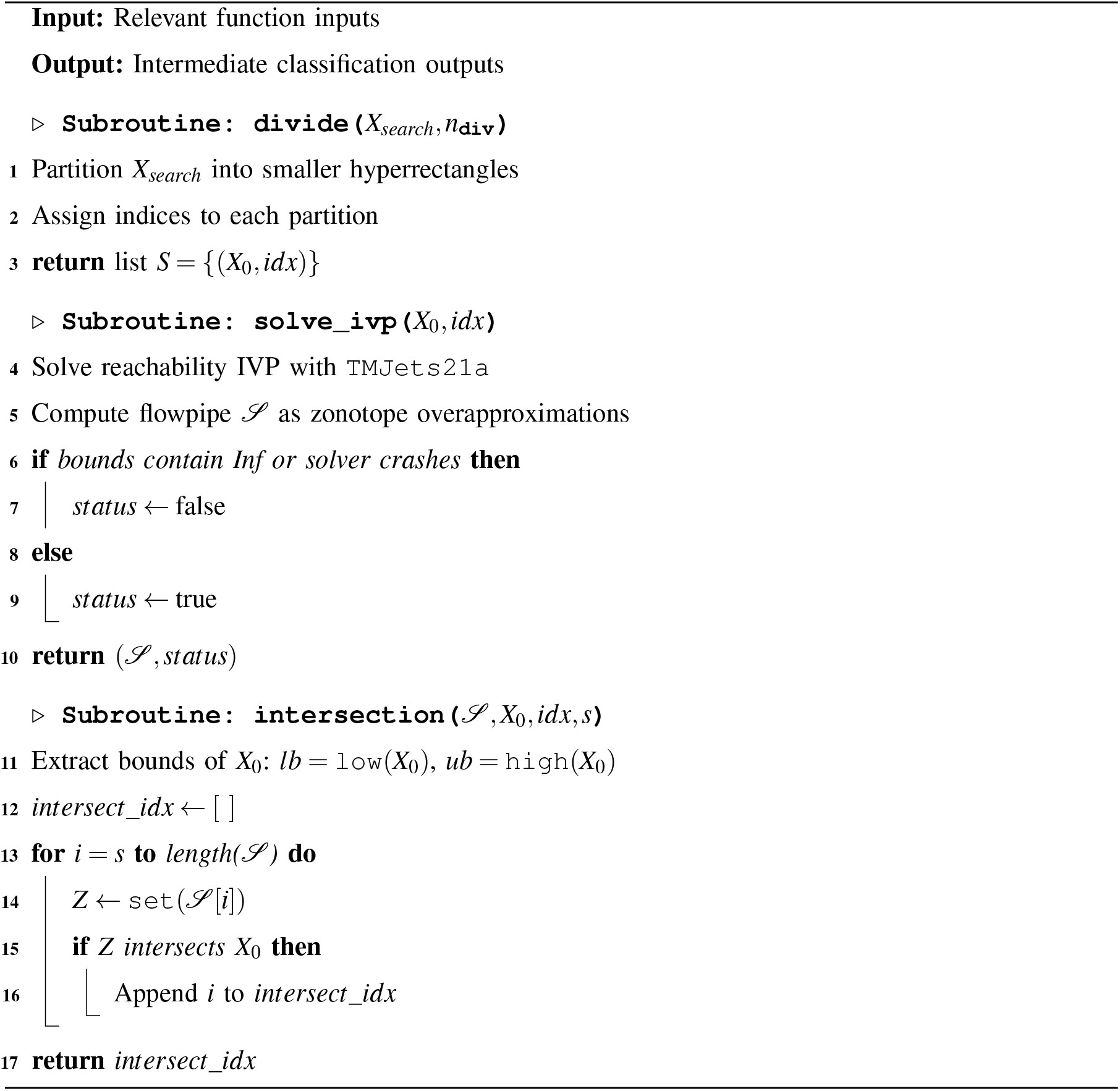

#### Algorithm 3: Subroutines for Reachability Analysis Based Localization of Limit Cycle (contd…)

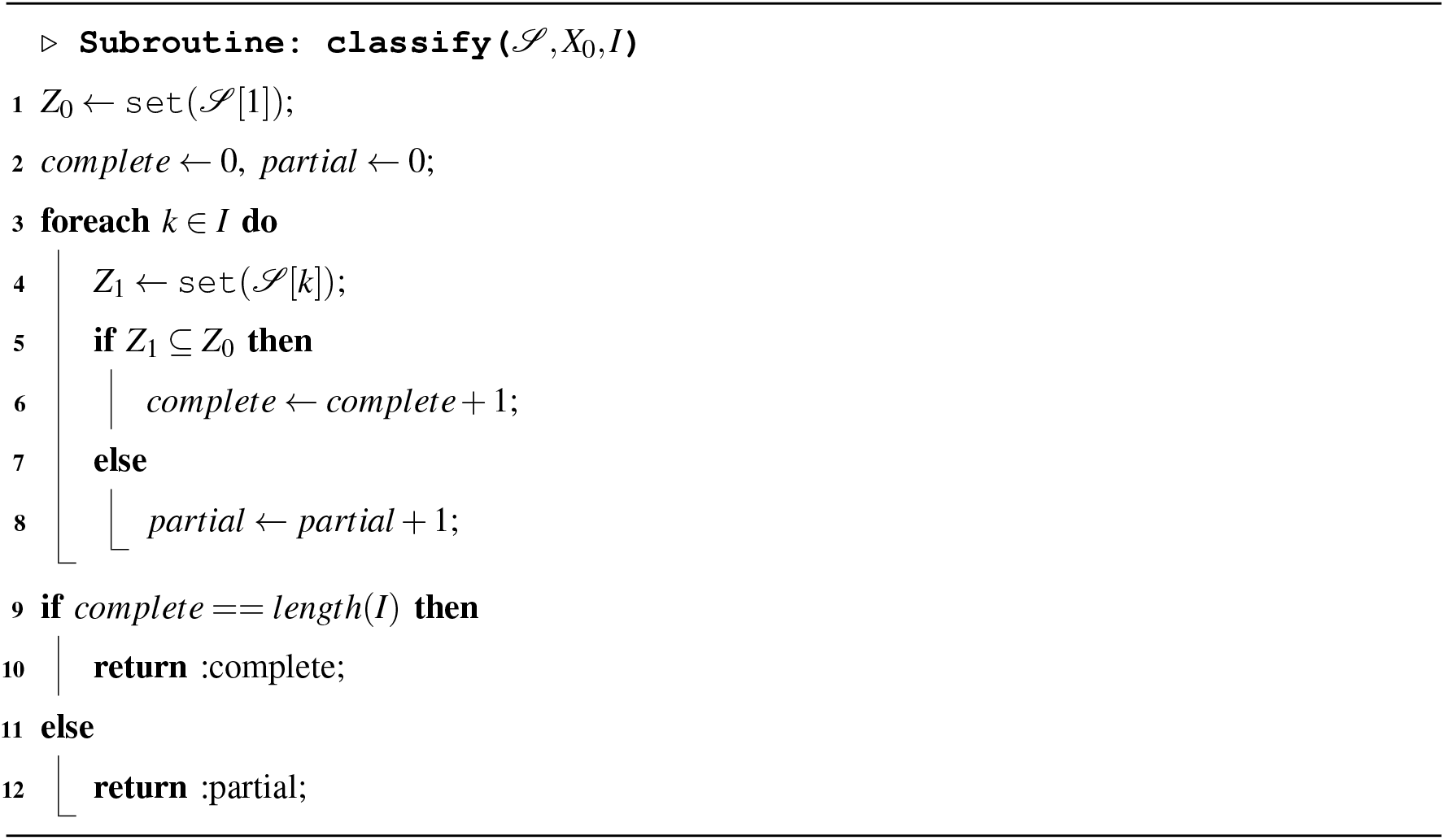

### C. Results and Discussions

The existence proof for the three-gene cyclic regulatory network within an invariant cube 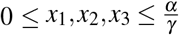, when divided into 2^3^ smaller boxes by the surfaces *x*_*i*_ = *x*_0_ (for *i* = 1, 2, 3), gives us six cross-sections of the smaller boxes that map onto themselves. We then illustrated the interval-based Reachability Analysis to localize periodic orbit through one of these cross-sections that map onto themselves in the three-gene cyclic regulatory network. The framework successfully divided the cross-section according to the underlying dynamical behaviour of the three-gene cyclic regulatory network. The methodology, based on partitioning the cross-section and analyzing the flowpipe of each partition, showed several findings, which are visually summarized in Fig. 1.

**Figure 1:**
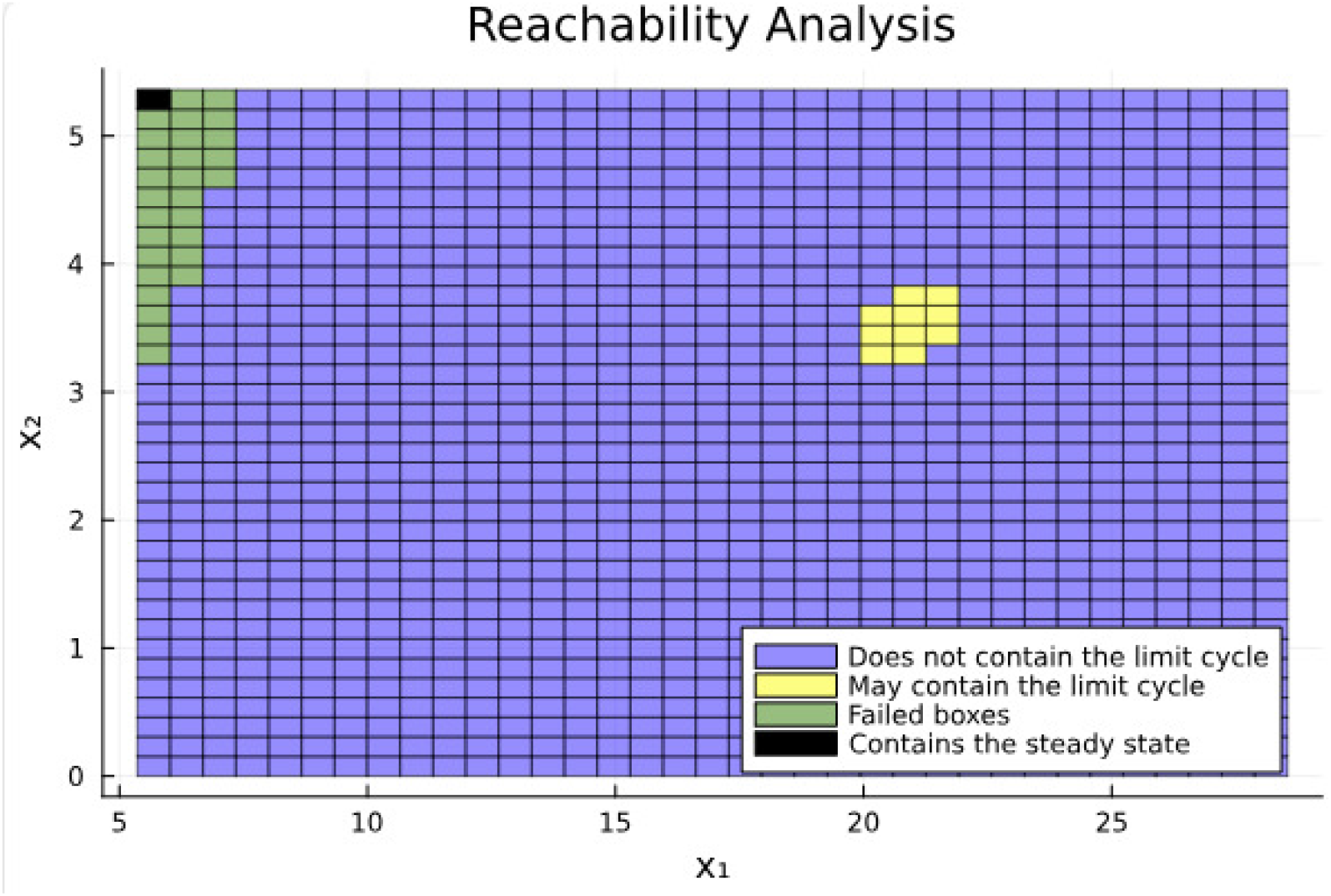
Projection of the cross-section, that maps onto itself, showing the geometric classification results. The black box contains the steady-state point and was excluded from the reachability analysis. The blue boxes is the region that does not contain the limit cycle. The cluster of yellow boxes signify the region that may contain the limit cycle. The green boxes are classified as the boxes that failed to return a solution for reachability analysis due to solver crash or blow up in the bounds due to overapproximation. The system parameters used in the simulation are: *α* = 26.26, *γ* = 0.92, *k* = 4, *n* = 5.

The analysis classified the vast majority of the partitioned region of this cross-section as not containing a periodic orbit. The black box in Figure 1, contains the unstable steady state (*x*_0_, *x*_0_, *x*_0_) (the value of *x*_0_ for this simulation is 5.361) of the system and is excluded from the Reachability Analysis. A distinct cluster of initial sets, marked in yellow in Figure 1, was classified as “May contain the limit cycle”. A cluster of green boxes around the box containing the steady state is classified as “Failed boxes”. This is because of the failure of the reachability analysis to return a solution. This can be attributed to solver crash or blow up in the bounds due to overapproximation. Further in Fig. 2 the numerical solution of the three gene cyclic regulatory network model is plotted along with one of the cross-section that maps onto itself. Fig. 3 provides a magnified view of the trajectory passing through the region classified by the method as “May contain the limit cycle”. This finding validates the method’s efficacy in localizing the region in which the limit cycle exists.

**Figure 2:**
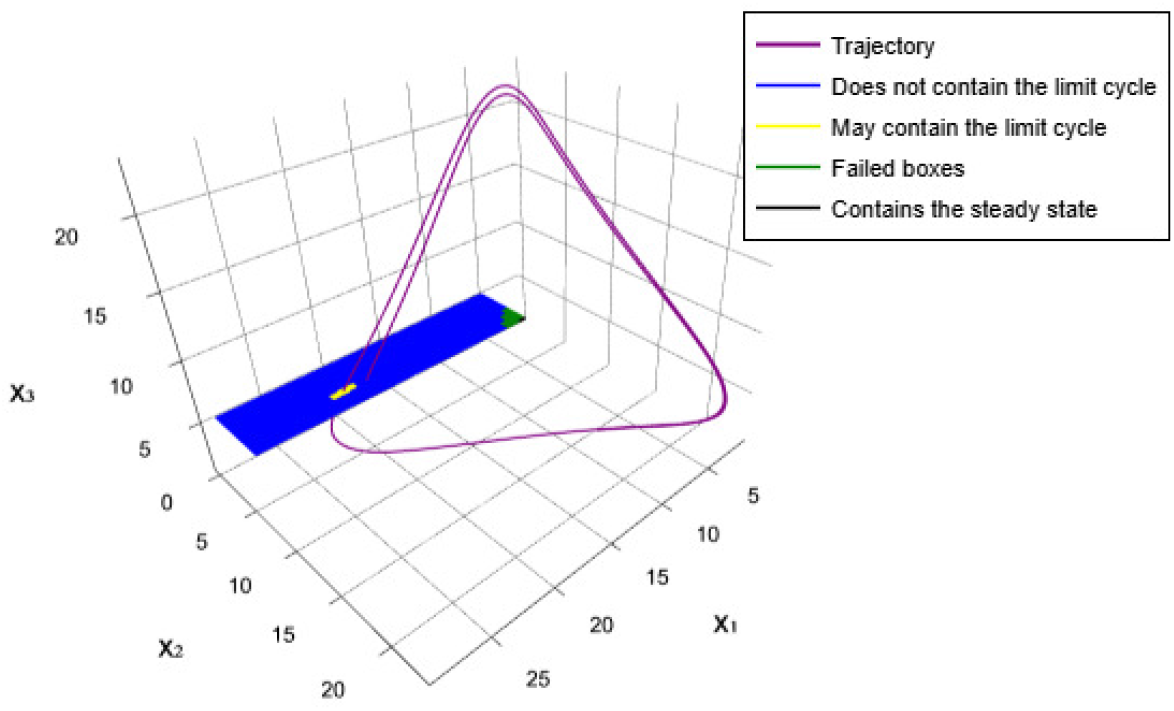
The numerical solution of the three-gene cyclic regulatory network with initial condition (19, 3.5, 5.361) passes through the region classified as “May contain the limit cycle” in Fig. 1. The system parameters used in the simulation are: *α* = 26.26, *γ* = 0.92, *k* = 4, *n* = 5. *x, y, z* axes represent the concentration of *i*−*th* (for *i* = 1, 2, 3) protein respectively.

**Figure 3:**
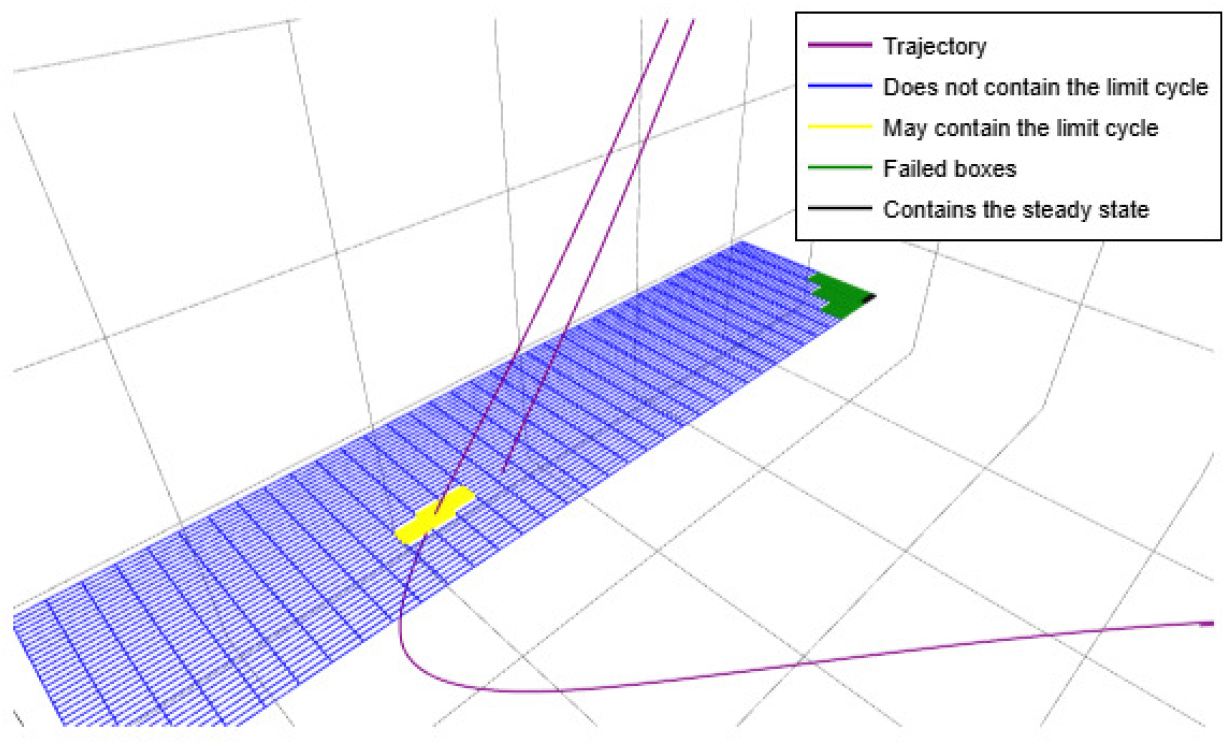
A magnified view of Fig. 2 showing the intersection of the numerical solution with the boxes classified as “May contain the limit cycle”.

## VI. Conclusion

We proved the existence of a limit cycle in a class of cyclic gene regulatory networks and provided a rigorous localization of this region of existence. The existence proof, based on Brouwer’s Fixed Point theorem, is an alternative and elementary proof for this class of systems, with the benefit of an enhanced geometric visualisation. This rigorous localization relies on an interval-based Reachability Analysis, which, in conjunction with Brouwer’s Fixed Point theorem, can provide a significantly tighter enclosure of the limit cycle trajectory.

For the existence proof, we constructed a positively invariant set for the network dynamics by exploiting the inherent boundedness and by creating a toroidal-like manifold region that eliminates hypervolume consisting of the unstable steady state and its associated stable manifolds. We found that a cross section of this manifold continuously maps onto itself under the system dynamics, showing via Brouwer’s Fixed Point theorem, the existence of a limit cycle. These results are illustrated using a 5-dimensional example. After establishing the existence of a limit cycle in cyclic gene regulatory networks, we employed interval-based Reachability Analysis to localize the region of its existence across the cross-sections that map onto themselves. The Reachability Analysis was demonstrated using a 3-dimensional example, offering a visual representation of the categorized regions.

## Footnotes

† The authors developed all algorithms presented in this manuscript. The LaTeX syntax and formatting were partially generated with the help of ChatGPT (OpenAI) as a support tool. After using this tool/service, the author(s) reviewed and edited the content as needed and take full responsibility for the content of the manuscript.

